# Eigenvector centrality mapping for ultrahigh resolution fMRI data of the human brain

**DOI:** 10.1101/494732

**Authors:** Gabriele Lohmann, Alexander Loktyushin, Johannes Stelzer, Klaus Scheffler

## Abstract

Eigenvector centrality mapping (ECM) is a popular technique for analyzing fMRI data of the human brain. It is used to obtain maps of functional hubs in networks of the brain in a manner similar to Google’s PageRank algorithm. ECM attributes a score to the time course of each voxel that reflects its centrality within the network. Voxels that are strongly correlated with many other voxels that are themselves strongly correlated with other voxels receive high scores. Currently, there exist two different implementations ECM, one of which is very fast but limited to one particular type of correlation metric whose interpretation can be problematic. The second implementation supports many different metrics, but it is computationally costly and requires a very large main memory. Here we propose two new implementations of the ECM approach that resolve these issues. The first is based on a new correlation metric that we call “ReLU correlation (RLC)”. The second method is based on matrix projections. We demonstrate the use of both techniques on standard fMRI data, as well as on high-resolution fMRI data acquired at 9.4 Tesla.

## 1 Introduction

Eigenvector centrality was originally proposed for investigating social behaviour [1, 2]. It later became the backbone of Google’s PageRank algorithm [3]. More recently, it was proposed as a tool for analyzing fMRI data of the human brain where it is used to obtain maps of functional hubs in networks of the brain [4]. Eigenvector centrality mapping (ECM) attributes a value to each voxel in the brain such that a voxel receives a large value if it is strongly correlated with many other voxels that are themselves central within the network. In the following, we briefly recapitulate ECM. For a more detailed description, see [4].

Let *S* be a symmetric *n* × *n* similarity matrix where entries *s*_*ij*_, *i, j* = 1,…*n* contain a pairwise similarity measure between time series in voxels *i* and *j*. The number of voxels *n* is determined by a user-defined mask which may cover the entire brain. In standard fMRI data with a spatial resolution of 3 × 3 × 3 mm, the number of brain voxels is around *n* ≈ 60, 000. However, in high-resolution images, this number can be much larger. For example, later on in this paper we will describe data acquired at 9.4T where *n* ≈ 500, 000.

At the core of the ECM algorithm is the computation of the principal eigenvector of *S*. Specifically, let *v* be the principal eigenvector of *S* and λ the corresponding eigenvalue, i.e. *Sv* = λ*v*. The eigenvector centrality score *v*_*i*_ of voxel *i* is then defined as the *i*-th entry of *v*.

The ECM algorithm computes the principal eigenvector *v* of *S* using power iterations. Specifically, we first initialize *v*_1_ = {1*/n,*…, 1*/n*}, and then iterate until convergence:

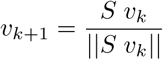

Generally, a few iterations suffice to arrive at the principal eigenvector *v*.

To be applicable for fMRI data analysis, it is necessary that the ECM algorithm yields unique results. However, eigenvalues and hence eigenvectors need not be unique. A way around this problem is to require that all entries of *S* must be non-negative, because the dominant eigenvalue λ of a irreducible matrix with non-negative entries has multiplicity 1 (Perron-Frobenius theorem [5]). For real symmetric matrices with non-negative entries, this means that the normalized corresponding eigenvector *v* is unique and has only non-negative entries.

The non-negativity requirement poses restrictions of the type of similarity metrics that can be used in this context. In particular, the Pearson correlation coefficient must be modified to meet this restriction.

One approach is to add the constant +1 to all correlation values. The advantage of this approach is that a very fast implementation becomes possible [6]. Specifically, let *X* be the *n* × *m* data matrix representing *n* voxels and *m* time points where each row of *X* represents a time course in one voxel normalized to 𝒩 (0, 1). Then the *n* × *n* correlation matrix is *XX*^*T*^ */n*. Let *E* be an *n* × *n* matrix with *e*_*i,j*_ = 1, ∀*i, j*. Power iteration can now be implemented very efficiently as follows.

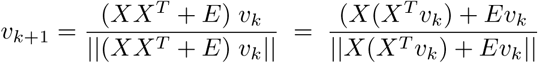

For ease of presentation, we will call this approach “ECM-ADD1”. As pointed out in [6], this implementation has the advantage that the full matrix *S* does not have to be kept in main memory at any time. Furthermore, the iterations are very fast. The disadvantage is that very small correlations are mapped to some intermediate values which may be hard to interpret. Also, the resulting ECM maps tend to have poor contrast.

The second approach is to set all negative correlations to zero and leave all positive correlations unchanged. Alternatively, one might use the absolute value of the correlation, or use some other metric that produces non-negative values such as mutual information or spectral coherence. See also [7–10] for more about measures of dependence. However, with such correlation metrics, the fast implementation approach shown above is not possible.

The implementation of the ECM algorithm as described in [4] supports various types of correlation metrics, and therefore requires that the matrix *S* is kept in main memory. This is generally prohibitive for high-resolution data. For example, if the number of voxels equals 500,000, then the memory requirement for *S* is around 465 GByte, see Fig. 1.

**Fig. 1:**
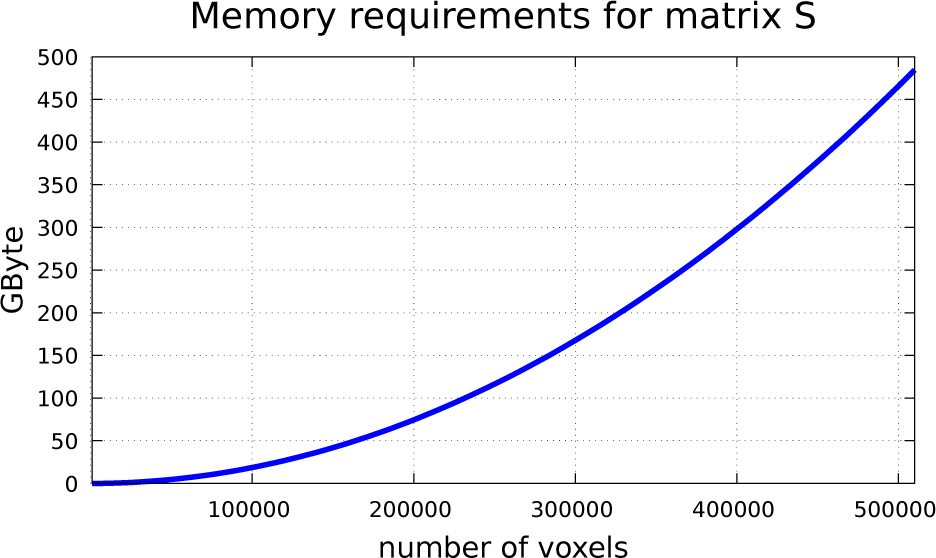
Memory requirements for storing matrix *S* in main memory. The size of the matrix S increases quadratically with the number of voxels. At 100,000 voxels, it is around 20 GByte. At 500,000 voxels, it is around 465 GByte. Here we assume that the matrix is represented using 4 Byte floating point values, and due to symmetry only one half of the matrix needs to be stored.

In this paper, we propose two methods for handling this problem. The first approach is to modify Pearson’s correlation coefficient such that it lends itself to a memory efficient and fast implementation. We call this modification “ReLU correlation (RLC)”. The second approach is based on matrix projections [11]. Here we use matrix projection as a technique for approximating the principal eigenvector in a memory efficient manner suitable for high-resolution data. This approach is not limited to any particular correlation metric. In the following, we describe both methods in more detail.

## 2 ECM based on ReLU correlations (RLC)

Let *X* be the *n* × *m* data matrix representing *n* voxels and *m* time points where each row of *X* represents a time course in one voxel normalized to zero mean and standard deviation of one. In order to ensure non-negativity, we propose to modify Pearson’s linear correlation by adding absolute values at each time point as follows.

### Definition

Let 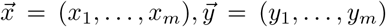 be two rows of *X* representing two voxels. We define the *ReLU correlation coefficient (RLC)* between 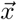 and 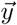 as:

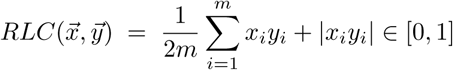

The RLC coefficient is zero if all pairs *x*_*i*_, *y*_*i*_ have discordant signs in all time points. The maximum value is obtained if 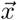 and 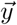 have a Pearson correlation of 1. Note that the RLC differs from Pearson’s correlation in the term |*x_i_y_i_*| that is added at each time point. The effect of this modification is that time points in which the signs of the two time series are concordant make a positive contribution to RLC, while all others are set to zero. This modification amounts to applying the Rectified Linear Unit (ReLU) filter at each time point. The ReLU filter is widely used in the context of artificial neural networks [12].

To illustrate the effect of this new metric, we randomly generated pairs of data with a predefined Pearson correlation *r* ∈ [−1, 1] and applied the RLC metric as well as the alternative non-negative metric (*r* + 1)/2 that is currently used in [6], see Figure 2. Note that the RLC metric reduces negative values of *r* to much smaller values than (*r* + 1)/2, and enhances positive values of *r*.

**Fig. 2:**
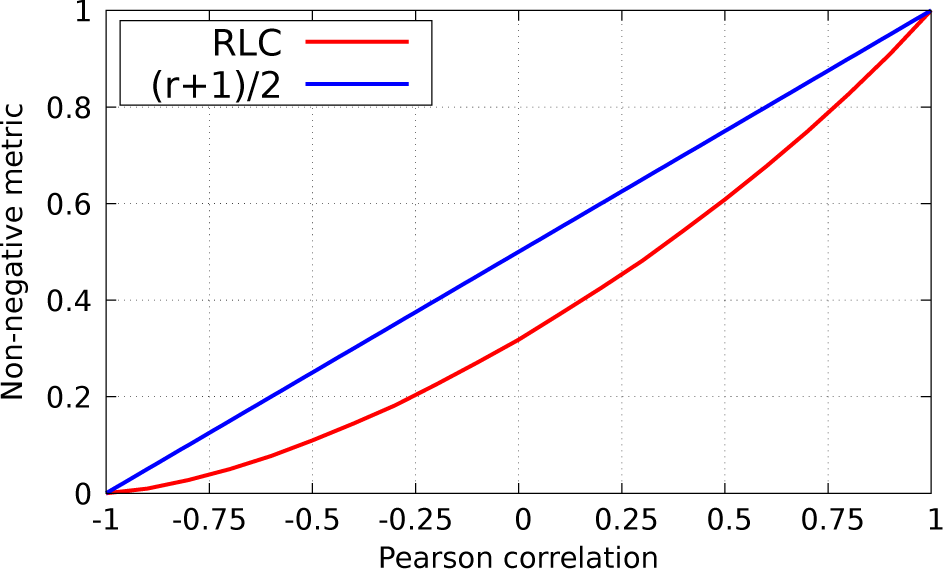
RLC versus Pearson correlation. We randomly generated set of data pairs with a given correlation coefficient of r ∈ [−1, 1]. The plot shows the RLC coefficient (red). The blue curve shows (r + 1)/2 which is the alternative approach that allows a fast implementation. Note that at r = 0 the RLC is mapped to rlc = 0.318, whereas (r + 1)/2 = 0.5. The data pairs (x_i_, z_i_), i = 1,…, n = 500000 were generated using 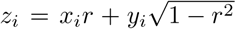 with x_i_, y_i_ randomly sampled from a standard Gaussian distribution.

The RLC correlation matrix can be computed via a single matrix multiplication by adding *n* columns to the data matrix *X*. More precisely, for all voxels *i* ∈ {1,… *n*}, we define

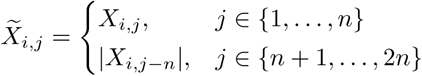

Thus, the RLC matrix of 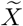 can be computed as

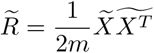

This formulation allows for a very efficient implementation of ECM. We first initialize *v*_1_ = {1/2*n,*…, 1/2*n*}, and then iterate until convergence

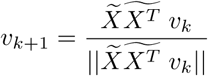

The term 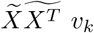 can be computed by first applying 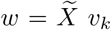, and then 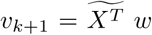. The advantage of this procedure is that the matrix 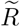 does not have to be kept in main memory. This makes this method applicable for high-resolution data in which the number of voxels can be extremely large so that the matrix 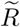 would require an excessive amount of main memory. A pseudocode is listed below.

#### Algorithm 1: ECM-RLC

1. Add *m* columns of absolute values to the data matrix: 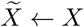
2. Initialize *v*_1_ = {1/2*n*,…, 1/2*n*}
3. Until convergence, repeat:

a. 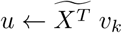
b. 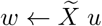
c. *v*_*k*+1_ ← *w*/||*w*||
4. The principal eigenvector is *v*_*k*+1_

In our implementation, convergence was established when ||*v*_*k*+1_ − *v*_*k*_|| < *ε* where *ε* is some small number (*ε* = 0.00001). By construction, the output vector *v*_*k*+1_ is normalized, i.e. ||*v*_*k*+1_|| = 1. Thus, the individual components of this vector have very small values. For better visualization, we therefore apply a componentwise scaling factor of 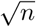 to the output vector as a final step.

## 3 ECM using matrix projection

In the following, we introduce a second approach for computing ECM. Here the emphasis is on reducing memory requirement so that ECM remains computable even with very large numbers of voxels. The main idea is to compute the approximation to principal eigenvector using matrix projections. Matrix projections were introduced by Halko et al. [11] as a technique for computing singular value decompositions.

As before, let *X* be the *n* × *m* data matrix representing *n* voxels and *m* time points where each row of *X* represents a time course in one voxel normalized to zero mean and standard deviation one. Let *S* denote a similarity matrix, and *S*^+^ denote a matrix where all negative entries are set to zero. The first step of the algorithm is to select a projection dimension *p* ≪ *n*. In our experiments, we used *p* = 32. The new ECM algorithm then consists of the following steps.

### Algorithm 2: ECM-project

1. Select a projection dimension *p* ≪ *n*, e.g. *p* = 32.
2. Generate a random matrix Ω ∈ ℝ^*m*×*p*^ sampled from a normal distribution 𝒩 (0, 1)
3. Compute *Y* = *S*^+^Ω
4. Perform SVD of *Y*, i.e. *Y* = *QΣR*^*T*^. *Q* forms an orthonormal basis for a range of *Y*.
5. Compute *E* = *Q*^*T*^ *S*^+^
6. Perform SVD of *E*, i.e. *E* = *U* Λ*W ^T^*
7. Compute *U* = *QU*
8. The first column of *U* contains the principal eigenvector.

Note that this algorithm can be implemented such that the similarity matrix *S*^+^ does not have to reside in main memory at any time. Instead, the partial matrix multiplications in steps 3 and 5 are performed in chunked block-wise fashion, and positive thresholding is applied to a partial matrix product result. Please note that steps 3 and 5 are computationally expensive. Computation time can be significantly reduced by summing only over a subset of voxels. This is possible because the matrix *S*^+^ is highly rank-deficient since the number of time points is in general much lower than the number of voxels. Therefore, low-rank approximations can be obtained without major loss in accuracy, see also [13] [14]. Alternatively, block matrix matrix multiplications can be very efficiently performed on GPUs, reducing the computation time even in high-resolution data to few minutes, see [15]. It is worth of note here that the same partial matrix matrix multiplication trick followed by thresholding can be applied for a power iteration method described above. The two singular value decompositions (steps 4 and 6) are performed on small matrices and very fast.

## 4 Experiments

### 4.1 Midnight scan club data (3T)

We applied the ECM-project algorithm to rs-fMRI data acquired by the “Midnight Scan Club” [16]. The Midnight Scan Club (MSC) data set contains repeated measurements of ten subjects (24-34 yrs; 5F). All MRI data were collected after midnight to control for time of day effects. Here we used the resting state fMRI data of subject 01 (run 01). The spatial resolution was (4*mm*)^3^, TR=2.2 seconds. 49 volumes were acquired producing a measurement time of 108 seconds. This data set was preprocessed using the toolbox “fMRIPprep 1.1.4.” [17]. It included a coregistration to the MNI template, and corrections for motion, and slice timing. We additionally applied a high pass filter which allowed frequencies faster than 1/100 Hz to pass, and a spatial Gaussian smoothing filter with fwhm=6mm. We constructed a region-of-interest mask covering the entire brain containing 28939 voxels. We applied ECM using a correlation metric in which negative Pearson correlation values were set to zero, and positive values were left unchanged. The results obtained with and without projection are shown in Figure 3. We compared the two results by computing a voxelwise relative difference *d*, where *d* is defined as *d* = (*u − v*)/*u* with *u, v* the ECM scores in a voxel. Note that |*d*| *<* 0.06 for all voxels. The histogram of both ECM results ranged in [0, 2].

**Fig. 3:**
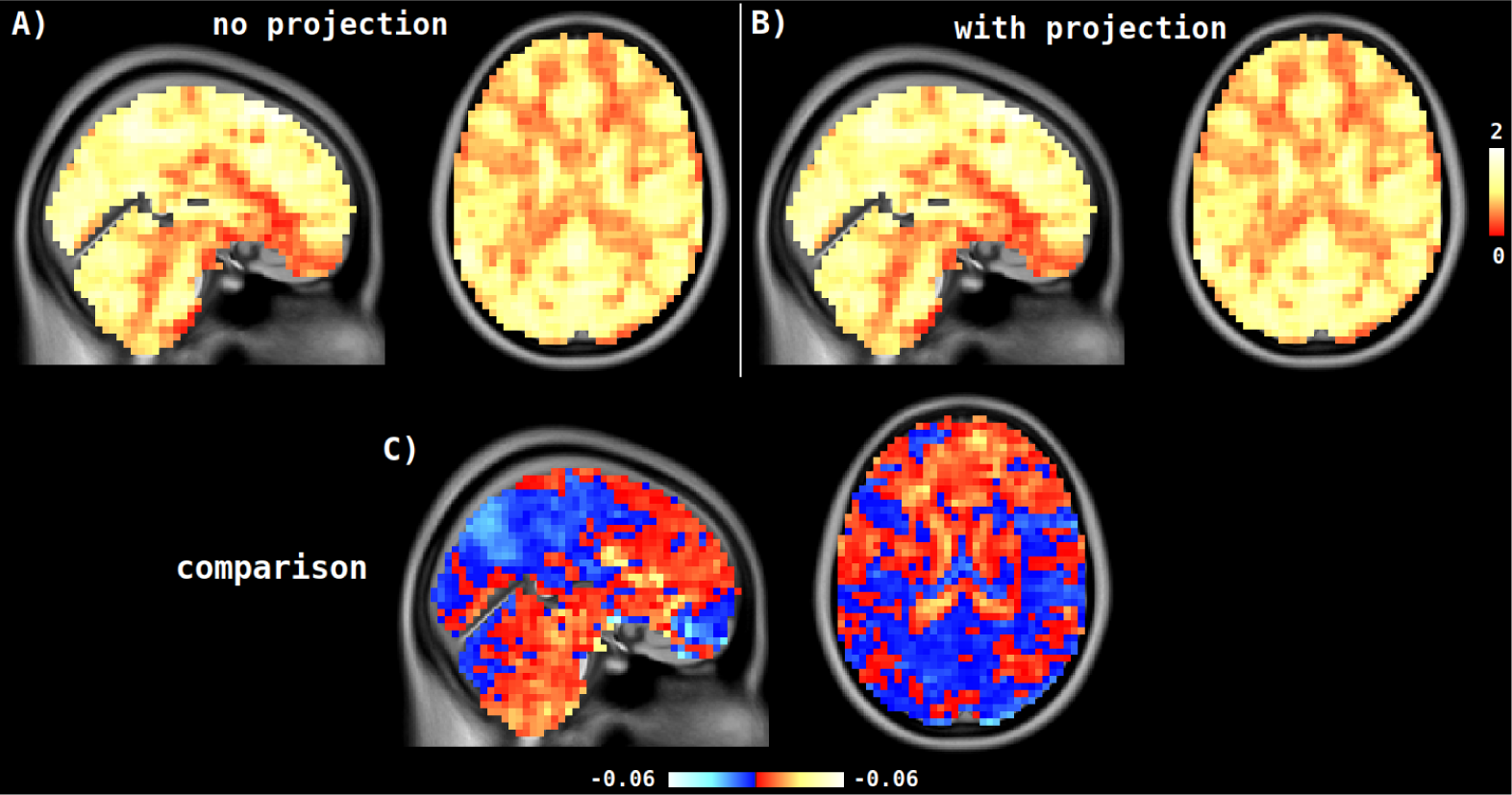
Comparing results of ECM with and without projection. The plot shows the results of ECM applied to the MSC-data using a correlation metric in which negative correlation values were set to zero, and positive values were left unchanged. The projection dimension was set to 32. Note that the relative difference did not exceed 0.06.

### 4.2 Ultrahigh resolution data (9.4T)

Resting state fMRI data of a single subject were acquired at a 9.4 Tesla Siemens Magnetom scanner. During the measurement time of 11.1 minutes, 330 volumes were acquired using TR=2.03 secs, TE=19ms, flipangle=70. Most of the brain was covered with 55 slices at a spatial resolution of (1.2*mm*)^3^, see Figure 4A. The subject (female, 29 yrs) was instructed to keep her eyes open. She gave written informed consent prior to the experiment, and was paid for her attendance. The study was approved by the local Ethics committee.

**Fig. 4:**
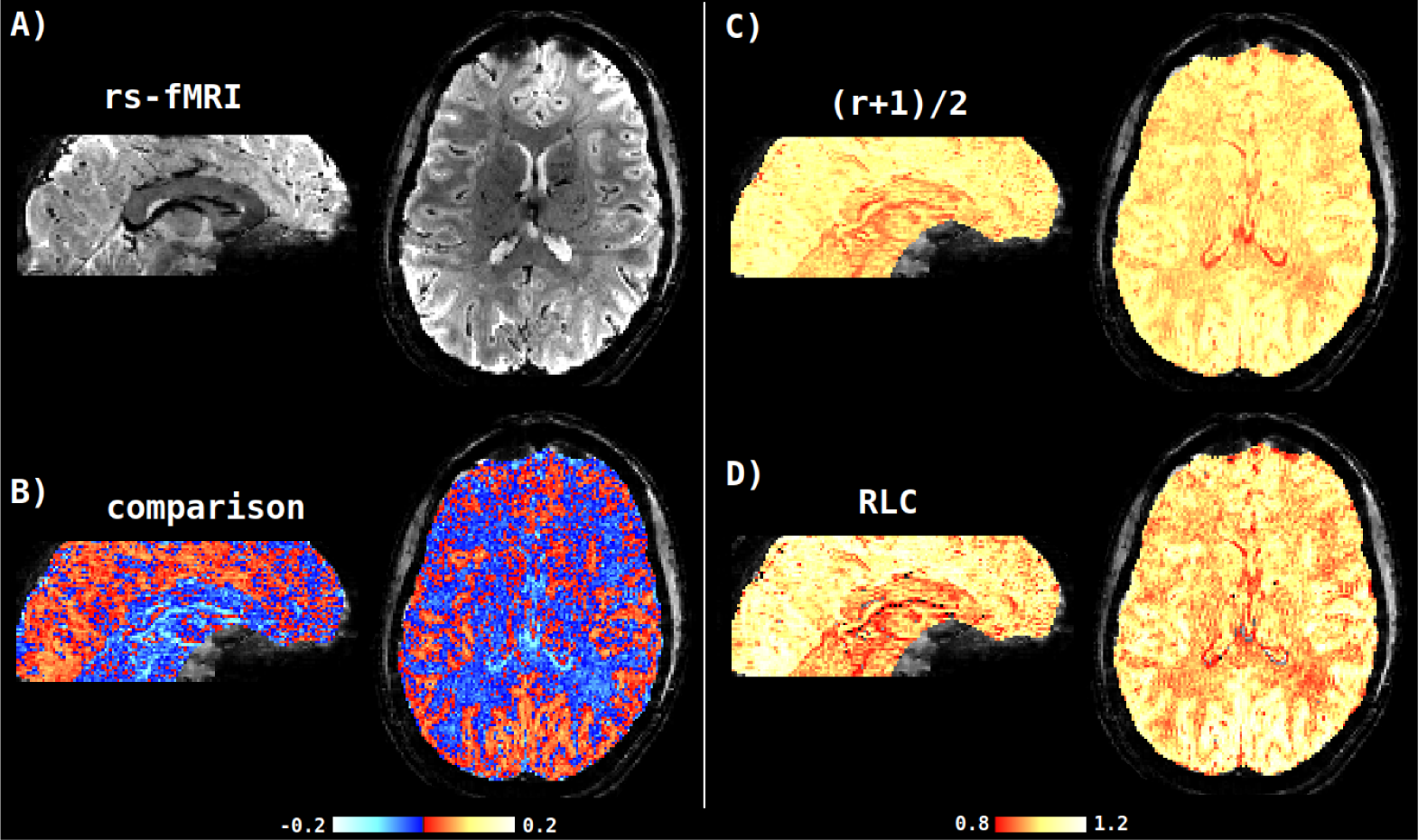
ECM-RLC applied to rs-fMRI data acquired at 9.4 Tesla. The plot shows the results of ECM-RLC (D) and previous metric ECM-ADD1 (C) applied to rs-fMRI data. The ROI mask contained 466462 voxels so that it is impracticable to store the entire correlation matrix in main memory. The comparison with ECM-ADD1 shows that ECM-RLC provides higher contrast particularly in grey matter regions (B). The computation time was around 20 seconds for both methods.

The data were corrected for motion and the baseline drift was removed using a highpass filter with a cutoff frequency of 1/100 Hz. We manually constructed a region-of-interest mask that contained 466462 voxels. The corresponding correlation matrix would require about 405 GByte of main memory making it impracticably large for standard desktop computers. We applied ECM-RLC as described above and ECM-ADD1 as in [6]. We compared the two approaches by computing a voxelwise relative difference as defined above. We found that ECM-RLC provides markedly better contrast particularly in grey matter regions.

## 5 Discussion

We have introduced two new algorithms for computing ECM. Both algorithms are memory efficient so that they can be used on high resolution data.

The first algorithm called “ECM-RLC” is based on a novel correlation metric called “RLC”. It allows for a very fast implementation comparable in speed only to the previous ECM-ADD1 algorithm. However, compared to ECM-ADD1 it showed markedly better contrast particularly in grey matter regions. This advantage may be explained by the fact that the ADD1 metric maps zero correlations to an intermediate value of 0.5, which does not well reflect the absence of a correlation. In contrast, the RLC metric maps zero correlations to a much lower value of around 0.31, see Figure 2. Thus, the RLC metric offers an advantage in terms of interpretablility compared to ADD1. Another advantage of ECM-RLC is that the ReLU filter is applied at each time point separately. This may help to better reflect dynamic changes in connectivity.

The second algorithm called “ECM-project” is computationally much more expensive. Its advantage compared to ECM-RLC is that it is not restricted to any particular type of correlation metric. ECM-project should be used whenever the RLC metric is not suitable and the number of voxels is so large that the classical implementation of ECM is not feasible due to memory limitations. Fast implementations of this algorithm are possible, especially when interpretability hardware is available. Our experiments show that the approximations obtained by ECM-project are highly accurate, even for very small values of the projection parameter *p*.

## Software availability

The software is available at: github.com/lipsia-fmri/lipsia.

## Acknowledgments

This work was partly funded by the European Commission H2020-GA-634541 CDS-QUAMRI.

## References

[1] Bonacich, P. Factoring and weighting approaches to clique identification. Journal of Mathematical Sociology 2, 113–120 (1972).

[2] Bonacich, P. Some unique properties of eigenvector centrality. Social Networks 29, 555–564 (2007).

[3] Langville, A. & Meyer, C. Google’s PageRank and Beyond: The Science of Search Engine Rankings (Princeton University Press, 2006). ISBN 0-691-12202-4.

[4] Lohmann, G. et al. Eigenvector centrality mapping for analyzing connectivity patterns in fMRI data of the human brain. PloS ONE 5, e10232 (2010).

[5] Frobenius, G. Ueber matrizen aus nicht negativen elementen. Sitzungsberichte der Königlich Preussischen Akademie der Wissenschaften 456477 (1912).

[6] Wink, A., de Munck, J., van der Werf, Y., van den Heuvel, O. & Barkhof, F. Fast eigenvector centrality mapping of voxel-wise connectivity in functional magnetic resonance imaging: implementation, validation, and interpretation. Brain Connectivity 2, 265–274 (2012). doi: 10.1089/brain.2012.0087.

[7] Reshef, D. Detecting novel association in large data sets. Science 334, 1518–1524 (2011).

[8] Lopez-Paz, D., Hennig, P. & Schölkopf, B. The randomized dependence coefficient. In NIPS (2013).

[9] Ding, A. & Li, Y. Copula correlation: an equitable dependence measure and extension of Pearson’s correlation. *arXiv* (2015). 1312.7214v4.

[10] Szekely, G., Rizzo, M. & Bakiriv, N. Measuring and testing the dependence by correlation of distances. The Annals of Statistics 35, 2769–2794 (2007).

[11] Halko, N., Martinsson, P. & Tropp, J. Finding structure with randomness: Probabilistic algorithms for constructing approximate matrix decompositions. SIAM Review 53, 217–288 (2011). https://arxiv.org/abs/0909.4061.

[12] Nair, V. & Hinton, G. Rectified Linear Units improve Restricted Boltzmann Machines. In Int. Conf. on Machine Learning (ICML) (Haifa, Israel, 2010).

[13] Xu, F., Yu, W. & Li, Y. Faster matrix completion using randomized svd. *arXiv* (2018). https://arxiv.org/pdf/1810.06860.pdf.

[14] Candès, E. & Plan, Y. Matrix completion with noise. Proceedings of the IEEE 98, 925–936 (2010).

[15] Fatahalian, K., Sugerman, J. & Hanrahan, P. Understanding the efficiency of GPU algorithms for matrix-matrix multiplication. In Proc. ACM SIGGRAPH/EUROGRAPHICS conf. on Graphics hardware (HWWS’04), 133–137 (ACM, 2004).

[16] Gordon, E. et al. Precision functional mapping of individual human brains. Neuron 95, 791–807 (2017).

[17] Esteban, O. et al. FMRIPrep: a robust preprocessing pipeline for functional MRI. *bioRxiv* (2018). doi: https://doi.org/10.1101/306951.

